# Global dissemination of tet(X3) and tet(X6) among livestock-associated Acinetobacter is sporadic and mediated by highly diverse plasmidomes

**DOI:** 10.1101/2021.08.02.454847

**Authors:** Ying-Ying Cheng, Yang Liu, Yong Chen, Fu-Man Huang, Rong-Chang Chen, Yong-Hong Xiao, Kai Zhou

**Author notes:** Correspondence: Kai Zhou, Address: Dongmen North Road No. 1017, Shenzhen People’s Hospital, 518020 Shenzhen, China.

## Abstract

The emergence of plasmid-borne *tet*(X) genes mediated high-level resistance of tigecycline largely threatening its clinical effectiveness. Currently, the dissemination pattern of plasmid-borne *tet*(X) genes remains unclear. In this study, 684 fecal and environmental samples were collected at six livestock farms, and 15 *tet*(X)-positive *Acinetobacter* isolates were recovered, mainly including 9 *tet*(X3)- and 5 *tet*(X6)-positive *A. towneri* strains. A clonal dissemination of *tet*(X3)-positive *A. towneri* was detected in a swine farm, while the *tet*(X6)-positive *A. towneri* strains mainly sporadically disseminated in the same farm. A *tet*(X3)-carrying plasmid (pAT181) was self-transmissible from a tigecycline-susceptible *A. towneri* strain to *A. baumannii* ATCC17978, causing a 128-fold and 64-512-fold increase in the MIC values of tigecycline and the other tetracyclines, respectively. Worrisomely, pAT181 was stably maintained and increased the growth rate of ATCC17978. Further identification of *tet*(X)s in 10,680 *Acinetobacter* genomes retrieved from GenBank revealed that, *tet*(X3) (n=249) followed by *tet*(X5)-like (n=61) and *tet*(X6) (n=53) are the prevalent alleles mainly carried by four species, and most of them are livestock associated. Phylogenetic analysis showed that most of *tet*(X3)- and *tet*(X6)-positive isolates disseminate sporadically. The structures of *tet*(X3) and *tet*(X6) plasmidomes are highly diverse and no epidemic plasmids have emerged yet. However, cross-species and cross-region transmissions of *tet*(X3) might have been mediated by several plasmids in a small proportion of strains. Our study evidence that *tet*(X3) and *tet*(X6) currently disseminate sporadically in *Acinetobacter*. Continuous surveillance for *tet*(X)s in the context of One Health is necessary to prevent them from transmitting to humans.

**Importance:** Recently identified plasmid-borne *tet*(X) genes highly challenged the efficiency of tigecycline, a last resort antibiotic for severe infection. Currently, the dissemination pattern of plasmid-borne *tet*(X) genes remains unclear. In this study, we first identified plasmid-borne *tet*(X)-positive *Acinetobacter* spp. strains from fecal and environmental samples collected at six livestock farms. A clonal dissemination of *tet*(X3)-positive *A. towneri* was detected in a swine farm, while the *tet*(X6)-positive *A. towneri* strains mainly disseminated sporadically in the same farm. A *tet*(X3)-carrying plasmid was found self-transmissible resulting in enhanced tigecycline resistance and growth rate. Further exploring a global dataset of *tet*(X)-positive *Acinetobacter* genomes retried from GenBank revealed that most of *tet*(X3) and *tet*(X6)-positive isolates share highly distant relationship, and the structures of *tet*(X3) and *tet*(X6) plasmidomes are highly diverse. Our study evidence that *tet*(X3) and *tet*(X6) disseminate sporadically in *Acinetobacter* and continuous surveillance for *tet*(X)s in the context of One Health is necessary.

## Introduction

Tigecycline is used to treat a wide range of clinical infection caused by Gram-positive and Gram-negative bacteria with multidrug resistance (MDR). With the global dissemination of carbapenemases and MCRs in recent years, this broad-spectrum tetracycline-family antibiotic has been raised to be a last line treatment regimen in clinical settings (1–6). However, the recent discoveries of transferable tigecycline inactivation genes [*tet*(X)s] particularly threaten the clinical efficacy of tigecycline (7, 8).

The first flavin-dependent monooxygenase gene *tet*(X) was identified in Tn*4351* and Tn*4400* encoded on the chromosome of *Bacteroides fragilis* in 1990 (9). Subsequently, numerous chromosome-encoded and plasmid-mediated *tet*(X) alleles, *tet*(X1) to *tet*(X14), have been reported in various species originating from animals, humans and environments (10–12). These Tet(X) variants, except Tet(X1), exhibited different levels of activity against almost all tetracyclines, including the fourth generation tetracycline (eravacycline) approved by the Food and Drug Administration (FDA) in 2018 (4, 12, 13). Remarkably, the first findings of plasmid-borne *tet*(X3) and *tet*(X4) identified in livestock-associated *Acinetobacter baumannii* and *Escherichia coli* strains in 2019 (7), respectively, raise the concern of horizontal transfer of tigecycline resistance. Since then, additional *tet*(X) alleles have been reported to be plasmid-borne, including *tet*(X5) and *tet*(X6) and their variants. Epidemiological studies reveal that these novel *tet*(X) orthologs have mainly circulated in animals in China due to the heavy uses of tetracyclines in husbandry (8). However, plasmids are currently rarely reported to be the transmissible vectors of *tet*(X)s although an increasing number of plasmid-borne *tet*(X)s has been detected. In some pioneer studies, IS*CR2* is highlighted to be the key element facilitating the horizontal transfer of *tet*(X)s through circular intermediates (14–17). Therefore, the role of plasmids in the dissemination of *tet*(X)s remains obscure.

Surveillance studies show that the *tet*(X) alleles have been detected in over 16 bacterial species with *Acinetobacter* spp. to be the predominate one, and *tet*(X4) is the only allele primarily detected in *E. coli* with a low prevalence (7, 11, 17–20). The *tet*(X)-positive *Acinetobacter* spp. isolates are mainly recovered from dairy cows, chickens and pigs in China (16, 21), and plasmid-borne and/or chromosomal-encoded *tet*(X3) and *tet*(X6) are prevalent among *Acinetobacter* spp. strains isolated from both humans and animals (7, 16, 20, 22, 23). A surveillance at avian farms showed that 1.6-18.3% *Acinetobacter* spp. strains were *tet*(X)-positive among seven provinces in China (23). Another surveillance for tigecycline-resistant *Acinetobacter* spp. from 2015 to 2018 in 14 provinces and municipalities in China reported that 2.3-25.3% *tet*(X)-positive isolates from pig farms, migratory birds and human samples were identified in 9 provinces (20). Currently, *tet*(X5) is solitarily detected in an *A. baumannii* strain from humans (22). However, it is unclear how the plasmid-borne *tet*(X)s disseminate among *Acinetobacter* spp., i.e. vertical transfer (clonal dissemination), horizontal transfer, and sporadic dissemination.

In this study, a surveillance of *tet*(X)-positive *Acinetobacter* spp. recovered from livestock and their surrounding environmental sources was performed at six livestock farms locating in Zhejiang province in 2019. The epidemiological and genetic characterizations of *tet*(X)-positive isolates and *tet*(X)-harboring plasmids were dissected. We further comprehensively investigated the population structure and distribution of *tet*(X)-positive *Acinetobacter* strains identified in the public database, as well as the plasmidome of *tet*(X3) and *tet*(X6).

## Results

### *A. Towneri* was the prevalent species carrying *tet*(X) genes among *Acinetobacter* strains collected in this study

Two hundred and ninety-two strains were recovered from 534 stool samples and 150 environmental samples collected from 2 swine farms, 2 dairy farms and 2 sheep farms, including 215 strains of *Acinetobacter* spp. and 77 strains belonging to other species. PCR screens of *tet*(X)s identified 23 positive isolates (7.88%; 23/292), including 15 *Acinetobacter* spp. isolates (6.88%; 15/218), 3 *Myroides odoratimimus* isolates and 5 *Empedobacter stercoris* isolates (Table 1). The 23 *tet*(X)-positive strains were exclusively isolated from swine farms. Twenty strains were recovered from the fecal samples of swine farm 1, and the 3 *M. odoratimimus* strains were from the soil samples of swine farm 2.

**Table 1.**
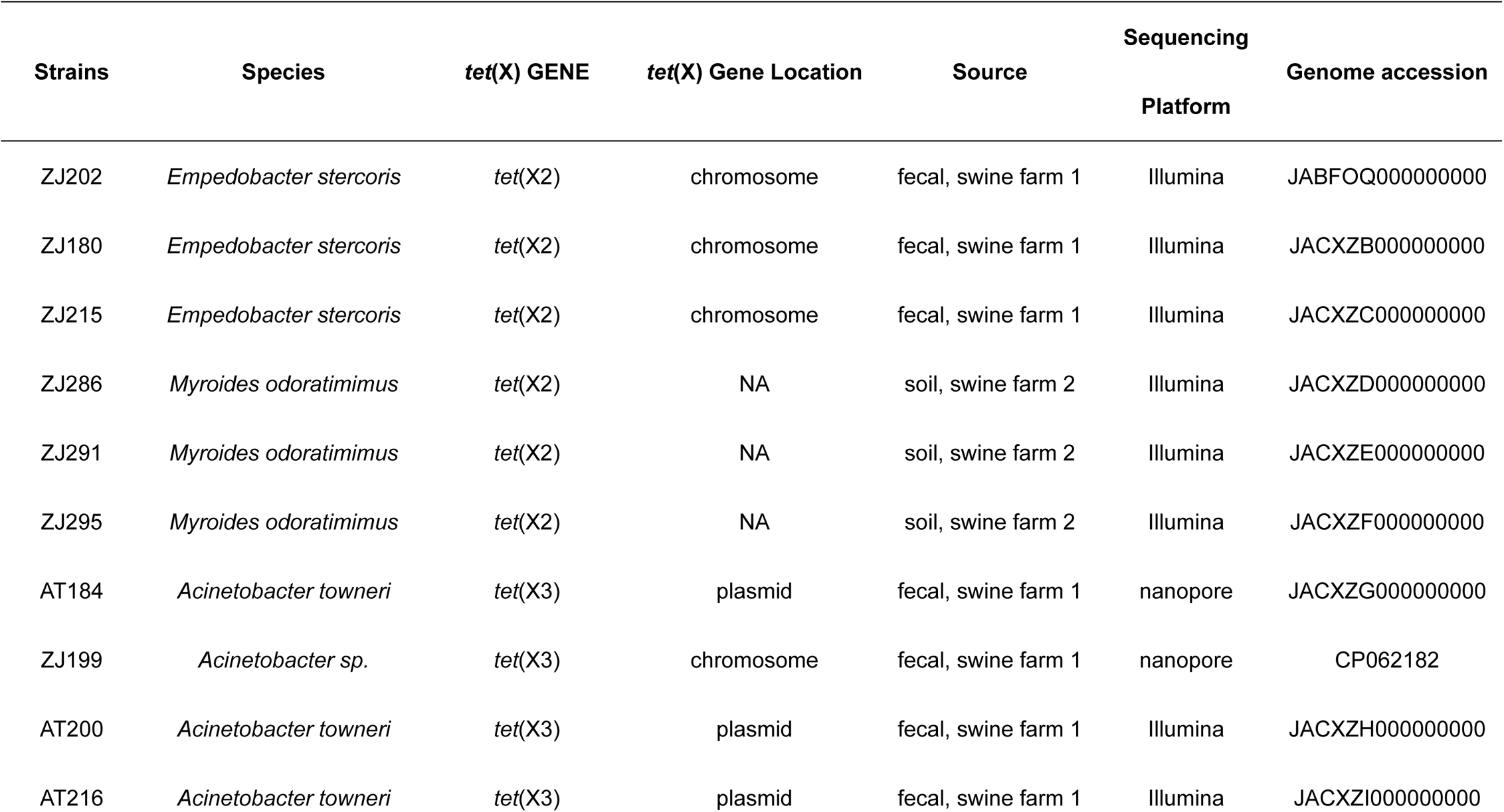
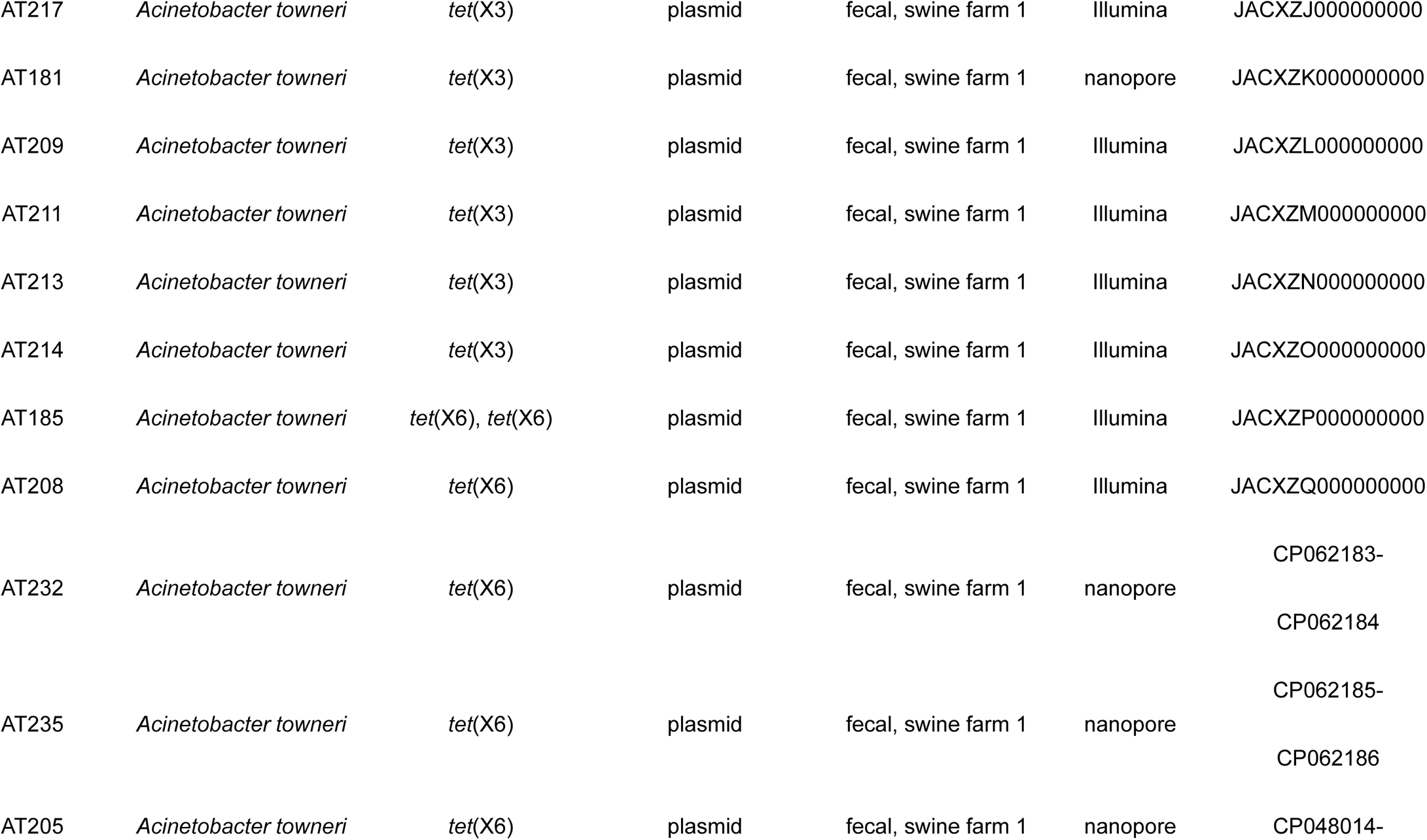
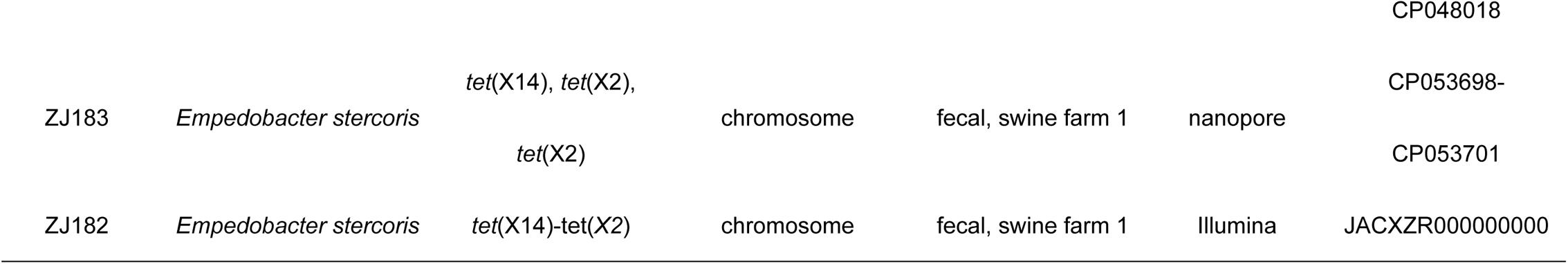
tet(X)-positive strains isolated in this study.

ANI analysis assigned the 15 *tet*(X)-positive *Acinetobacter* spp. isolates to *A. towneri* (n=14) and an unnamed species (n=1) (Table 1), suggesting that *A. towneri* was the prevalent species carrying *tet*(X)s in *Acinetobacter* spp. population circulating at swine farms. Four different *tet*(X) alleles were detected in the 23 isolates, including *tet*(X2) detected in 5 *E. stercoris* isolates and 3 *M. odoratimimus* isolates, *tet*(X3) in 9 *A. towneri* strains and 1 strain (ZJ199) belonging to the unnamed species, *tet*(X6) in 5 *A. towneri* strains, and *tet*(X14) in 2 *E. stercoris* strains (ES183 has been described previously (10)) (Table 1). One *A. towneri* strain (AT185) carried two copies of *tet*(X6). To our knowledge, this is the first report of two copies of *tet*(X6) identified in single strain. The phylogenetic analysis of 15 *tet*(X)-positive *Acinetobacter* spp. isolates showed that all but one *tet*(X3)-carrying *A. towneri* strains (8 out of 9) clustered together with 3-36 SNPs (Figure 1), suggesting a clonal dissemination of *tet*(X3) occurred in the swine farm. The other *tet*(X3)-carrying *A. towneri* strain AT200 clustered with the *tet*(X6)-carrying strains with 27,664-30,557 SNPs (Figure 1). All but two *tet*(X6)-positive strains showed distant relationship (26,876-31,071 SNPs), indicating that they disseminated sporadically.

**Figure 1.**
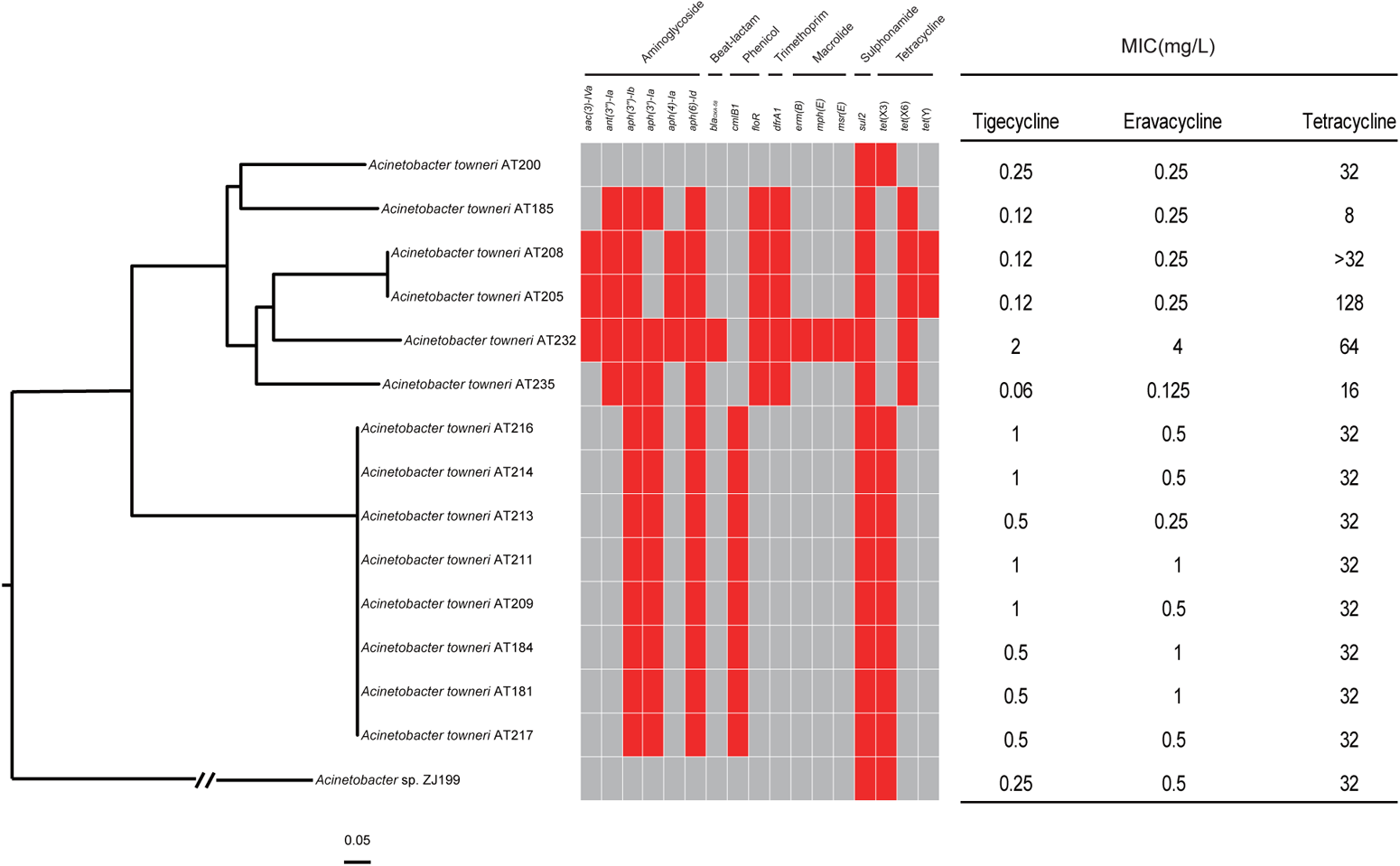
Phylogenetic analysis of *tet*(X)-positive *Acinetobacter* isolates collected in this study. The core-genome SNPs of *tet*(X)-encoding strains were used to generate the phylogenetic tree. The tree is rooted at strain ZJ199. The ARGs of each strain are exhibited by heatmap, and the existence of ARGs is in red. MIC values of each strain against tetracyclines are listed. AT205 has been reported previously [24].

### Antimicrobial resistance profile of *tet*(X)-carrying isolates

AST results showed that 34.78% (8/23) of *tet*(X)-positive isolates were resistant to tigecycline with MIC values at 1-2 mg/L, and the other 15 isolates showed MIC values at 0.06-0.5 mg/L (Table 2). These tigecycline resistant strains encompass 4 *tet*(X3)-positive *A. towneri* isolates, 1 *tet*(X6)-positive *A. towneri* isolate, 2 *tet*(X2)- and *tet*(X14)-positive *E. stercoris* isolates and 1 *tet*(X2)-positive *M. odoratimimus* isolate. Five tigecycline-resistant strains (3 *A. towneri* isolates and 2 *E. stercoris* isolates) additionally exhibited resistance to the newly FDA-approved eravacycline with MIC values at 1-4 mg/L. Except that the strain (AT185) carrying 2 copies of *tet*(X6) was susceptible to tetracycline, the other 14 *Acinetobacter* spp. strains were resistant to tetracycline with MIC values ≥ 16 mg/L (Table 2). Strain AT232 showed significantly higher resistance to tetracyclines than the other 13 strains, which might be caused by the presence of a two component system AdeSR involved in the expression of the AdeABC efflux pump (24). In addition, 26.7% (*n* = 4) and 13.3% (*n* = 2) *Acinetobacter* spp. strains showed resistance to ciprofloxacin and doxycycline, respectively (Table 2). All of *tet*(X)-positive *Acinetobacter* spp. isolates were susceptible to colistin and carbapenems. *M. odoratimimus* isolates were resistance to both colistin and carbapenems due to intrinsic resistance (25).

**Table 2.**
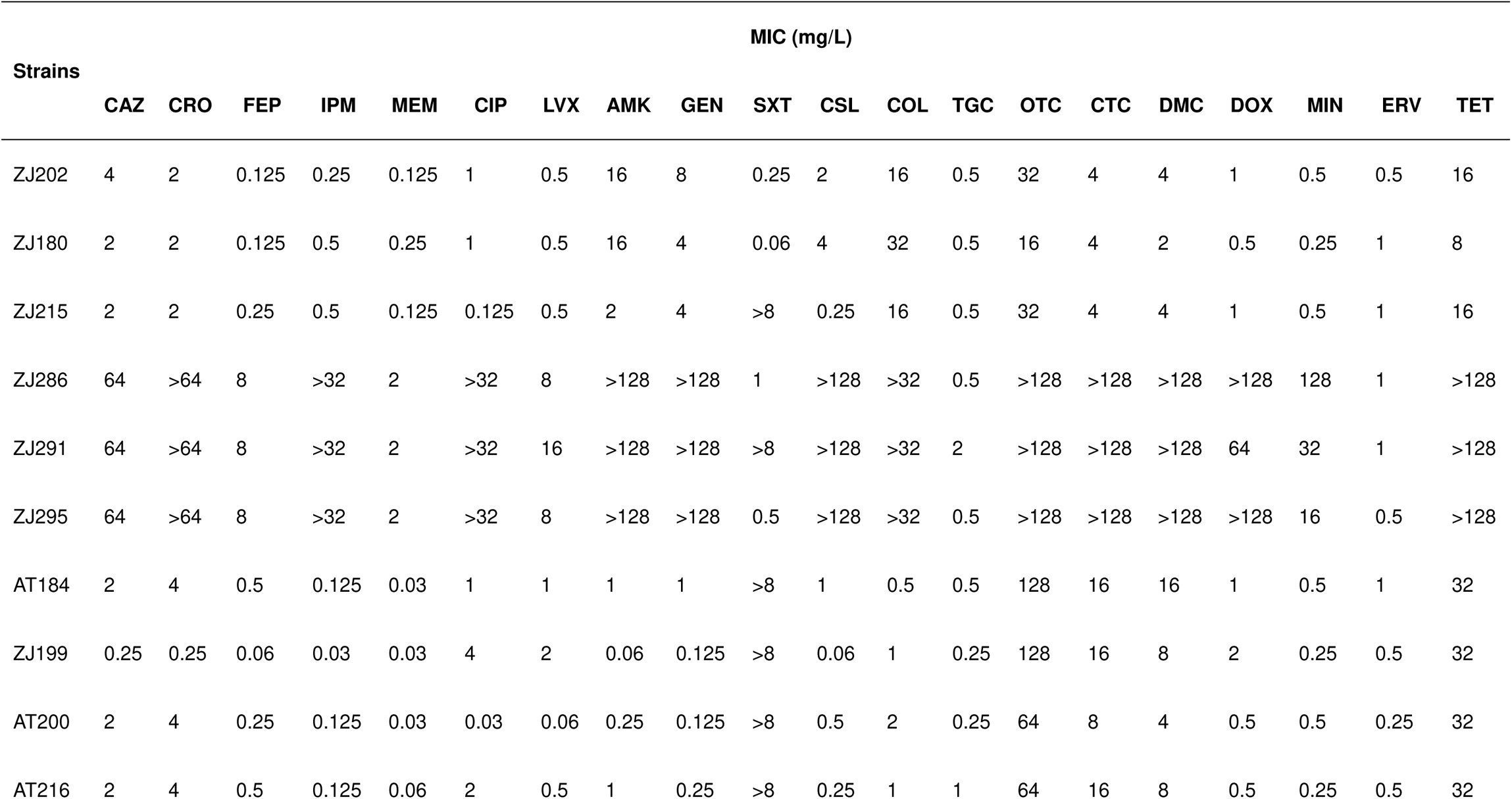
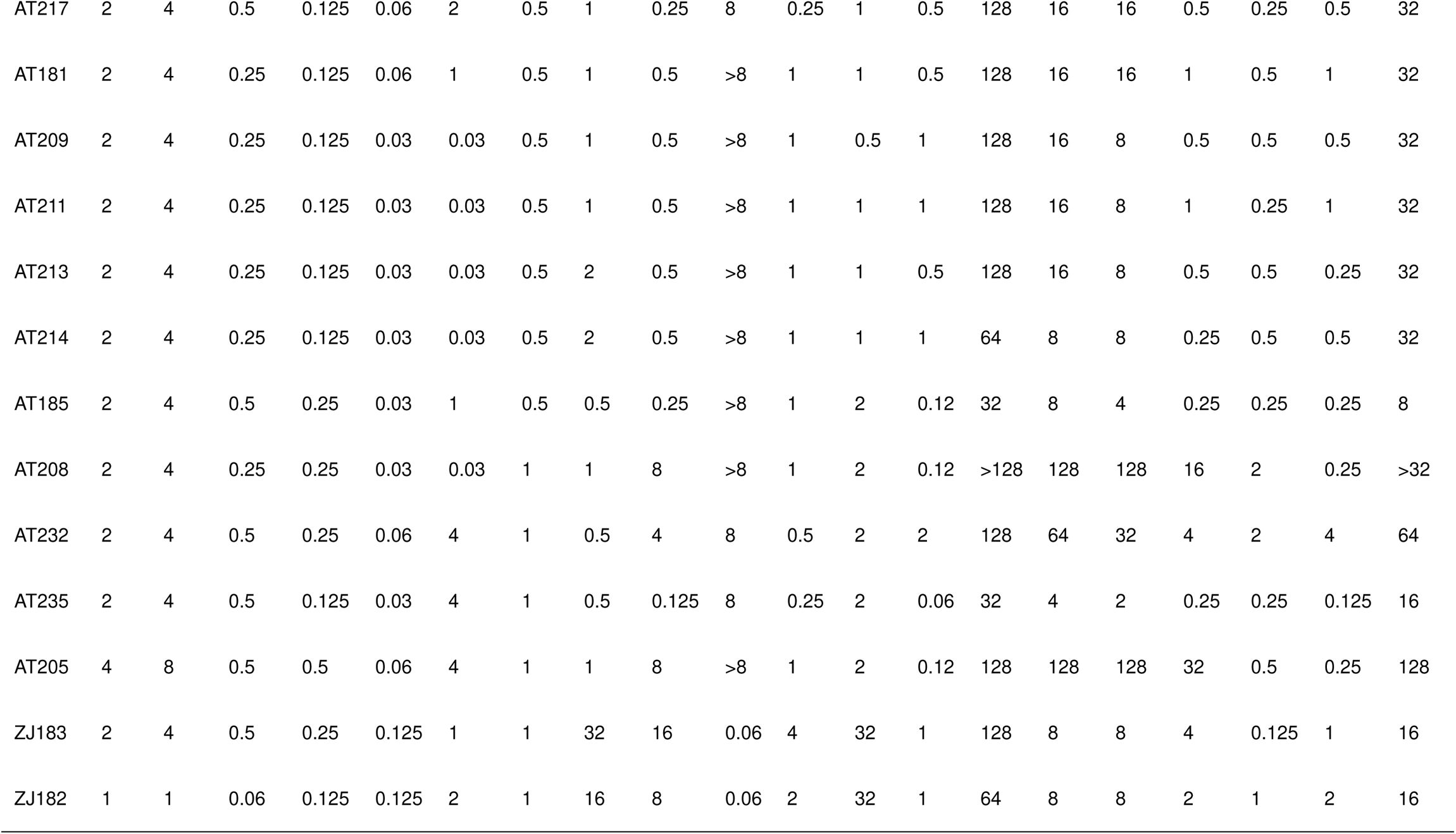
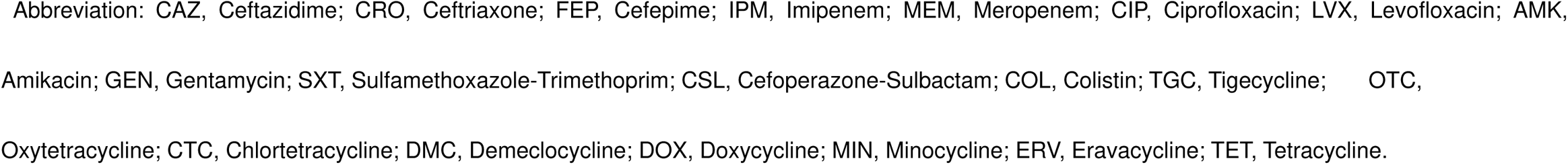
MIC values of antibiotics tested in this study.

*In silico* analysis of ARGs among *Acinetobacter* spp. strains showed that the number of ARGs detected in the strain ZJ199 [*sul2* and *tet*(X3)] was much less than that in *A. towneri* strains (Figure 1). All of *A. towneri* strains were MDR, and more ARGs were detected in the *tet*(X6)-carrying clone (mean=8.67; median=9) than in the *tet*(X3)-carrying clone (mean=6; median=6) albeit not significant (p > 0.05) (Figure 1). The 8 strains of the *tet*(X3)-carrying clone shared an identical resistome [*aph(3’’)-Ib*, *aph(3’)-Ia*, *aph(6)-Id, cmlB1*, *sul2* and *tet*(X3)], further supporting the clonal dissemination (Figure 1). While the resistome of the *tet*(X6)-carrying strains was highly diverse, including *aacC4*, *ant(3’’)-Ia* and *aph(4)-Ia* resistant to aminoglycoside; *bla*_OXA-58_ resistant to beta-lactam; *floR* resistant to phenicol; *dfrA1* resistant to trimethoprim; *erm(B)*, *mph(E)* and *msr(E)* resistant to macrolide; *tet*(X6) and *tet*(Y) resistant to tetracyclines (Figure 1). The resistome of *E. stercoris* and *M. odoratimimus* was different from that of *Acinetobacter* spp. (Table S1). *E. stercoris* strains carried *tet*(X2) and *tet*(X14) resistant to tetracyclines; *mef*(C) and *mph*(G) resistance to macrolide; and *bla*_EBR-1_ resistant to beta-lactam. *M. odoratimimus* strains carried *tet*(X2) and *tet*(36) resistant to tetracyclines; *ereD* resistant to macrolide; *bla*_MUS-1_ resistant to beta-lactam; and *sul2* resistant to macrolide (Table S1).

### *tet*(X3) and *tet*(X6) were harbored by various plasmids

To understand the vectors of the two prevalent *tet*(X) alleles, i.e. *tet*(X3) and *tet*(X6), the representative *tet*(X3)- and *tet*(X6)-carrying *Acinetobacter* spp. strains (AT181, AT184, and ZJ199; AT232 and AT235) were chosen additionally for long-read sequencing based on their antimicrobial resistance profiles and genetic environments of *tet*(X)s. The hybrid assembly confirmed that *tet*(X3) and *tet*(X6) were plasmid-borne in the four *A. towneri* strains, and a chromosome-encoded *tet*(X3) was detected in strain ZJ199.

The *tet*(X3)-carrying plasmids detected in AT181 (pAT181) and AT184 (pAT184) were identical with a size of 75,969-bp, and were circularized (confirmed by PCR). These two plasmids were untypable with an average GC content of 42.5%. Multiple ARG genes were carried by the two plasmids, including *aph(3’)-Ia*, *aph(3’’)-Ib*, *aph(6)-Id*, *sul2*, and *tet*(X3). Blast analysis of the nucleotide sequence of pAT181 in GenBank showed that the best match was a transferable *tet*(X3)-harboring plasmid p10FS3-1-3 (CP039146) (100% identity; 97% coverage) carried by a novel species of *Acinetobacter* (20). Other plasmids sharing a high similarity with pAT181 included a *tet*(X5)-harboring plasmid pAB17H194-1 (99.95% identity; 86% coverage) carried by an *A. pittii* strain and a *tet*(X3)-harboring plasmid p18TQ-X3 (CP045132, 99.99% identity; 80% coverage) carried by an *A. indicus* strain. These data suggest that pAT181-like plasmids have disseminated among various species of *Acinetobacter*.

pAT181 was used as a reference to perform blast comparisons among our *tet*(X3)-carrying strains to evaluate the genetic similarities of the other *tet*(X3)-carrying plasmids. The results revealed a conserved backbone shared by *tet*(X3)-carrying plasmids harbored in the 8 clonal strains with a coverage and nucleotide-acid identity >90% (Figure S1A). The *tet*(X3)-carrying plasmid carried by strain AT200 showed a different plasmid backbone with identity >90% and coverage <50% to pAT181 (Figure S1A). The best match of pAT200 was p10FS3-1-3 with 58.77% coverage and 70% identity, indicating that pAT200 might be a novel plasmid.

The two *tet*(X6)-harboring circularized plasmids pAT232 and pAT235 showed as low as 38% coverage and 99.95% identity between each other, suggesting that they were two different plasmids. pAT232 was 186,508-bp in length with GC content of 41.03%. Blasting in GenBank showed that the best matches of pAT232 were a *tet*(X6)-carrying plasmid pAT205 (CP048015) (76% coverage and 99.99% identity) carried by an *A. towneri* strain AT205 isolated in the same swine farm (26), and a *tet*(X)-negative plasmid p19110F47-2 (CP046044) (70% coverage; 99.99% identity) carried by an *A. towneri* strain isolated from the pig. pAT235 was 124,466-bp in length with GC content of 41.16%. The best matches of pAT235 were pAT205 (49% coverage; 100% identity) and a *tet*(X3)-harboring plasmid pGX7 (CP071772) (44% coverage and 99.95% identity) detected in an *A. towneri* strain isolated from the pig in China. These data suggest that pAT232 and pAT235 might originate from *A. towneri* associated with pigs.

When pAT232 was used as a reference to identify the plasmids of *tet*(X6) in the other *tet*(X6)-positive strains collected here, AT208 showed the highest similarity with pAT232 (77.84% coverage; 99.16% identity) (Figure S1B). When pAT235 was used as a reference, AT185 shared 100% coverage and 94.51% identity (Figure S1C), suggesting that a pAT235-like *tet*(X6)-encoding plasmid was harbored in AT185. Of note, AT185 was genetically distant from AT235 with 30,097 SNPs (Figure 1), suggesting that the horizontal transfer of pAT235-like plasmid might have occurred between the two strains. A pAT205-like *tet*(X6)-harboring plasmid was detected in AT208 when pAT205 was used as a reference (100% coverage; 96.48% identity) (Figure S1D). These results reveal that horizontal transfers of *tet*(X6)-carrying plasmids might have occurred in few strains.

### Genetic environment of *tet*(X3) and *tet*(X6)

The genetic environment of plasmid-borne *tet*(X3) [ΔIS*CR2*-*xerD*-*tet*(X3)-*res*-IS*CR2*] detected in the 8 of 9 *A. towneri* strains was identical, which was highly similar with that of the prototype detected in *A. baumannii* strain 34AB (7) (Figure 2A). To fully understand the distribution of this genetic environment among *tet*(X3)-carrying *Acinetobacter* strains, we blasted it against 249 *tet*(X3)-carrying *Acinetobacter* genomes retrieved from GenBank (see below), and results showed that 21.3% (53/249) genomes carry the fragment ΔIS*CR2*-*xerD*-*tet*(X3)-*res*-IS*CR2* locating on a single contig with >90% identity and >90% coverage. The proportion increased to 86.35% (215/249) when matches on different contigs were counted together, implying that this might be the major structure encoding *tet*(X3) in *Acinetobacter* spp.. A different genetic environment of *tet*(X3) [IS*4*-IS*4-tet*(X3)-*res*-ΔIS*CR2*] was detected on the chromosome of strain ZJ199, in which IS*CR2* and *xerD* located at the upstream of *tet*(X3) were replaced by two copies of IS*4* (Figure 2A). Inspection of the wider context of *tet*(X3) in strain ZJ199 showed that two copies of IS*4* adjacent to *sul2* and *glmM* located at the downstream of ΔIS*CR2* as found in an *A. indicus* strain AI2 (16) (Figure 2A). This results in a putative IS*4* bracketed transposon, which might be responsible for the mobilization of *tet*(X3) and *sul2*.

**Figure 2.**
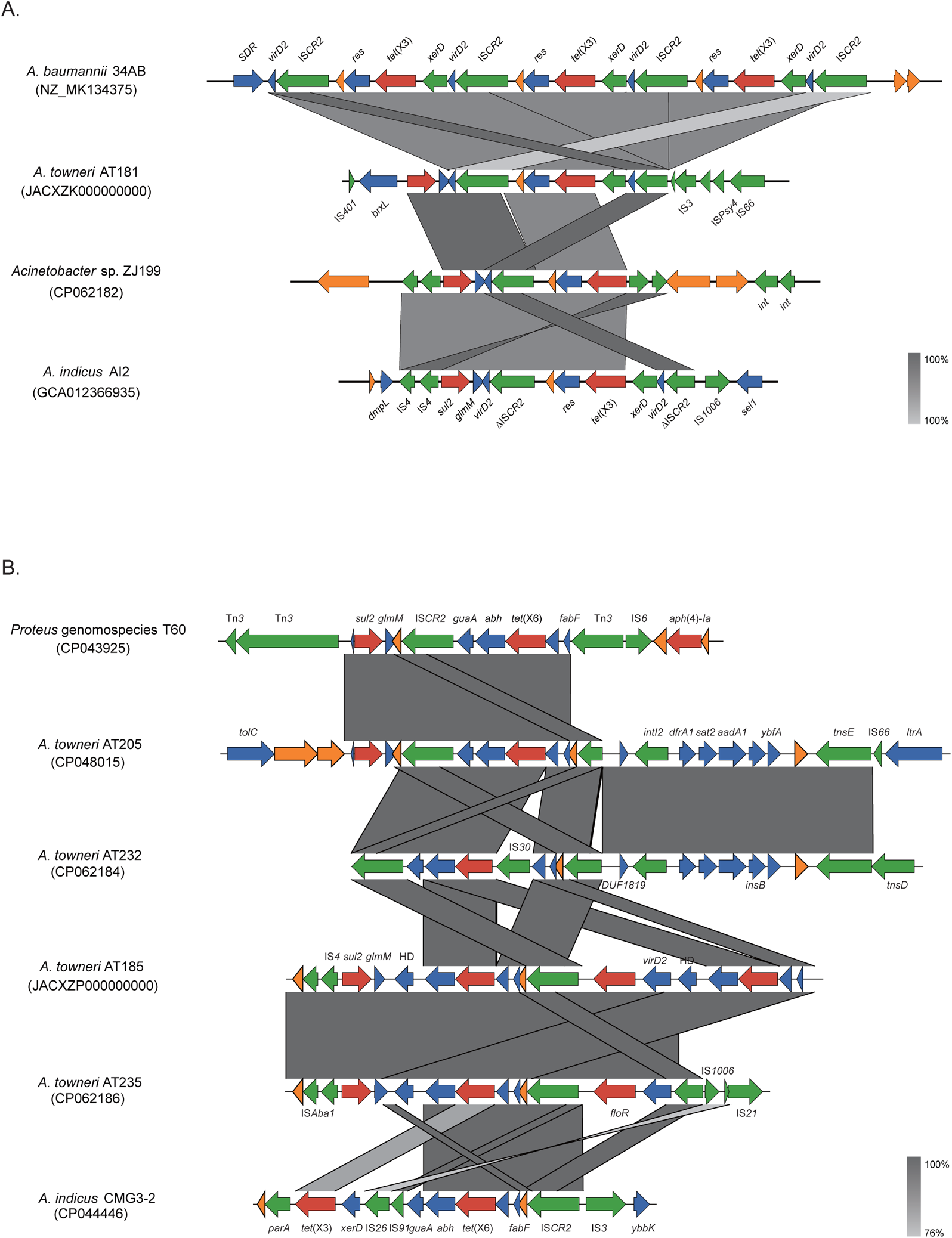
Comparison of genomic context of *tet*(X3) (A) and *tet*(X6) (B) identified in *Acinetobacter* spp. isolates. Genes are indicated by colour-coded arrows dependent on the functional annotations and direction of transcription. ARGs are in red; mobile genetic element genes are in green; other function genes are in blue; hypothetical genes are in orange. The **g**enomic context of *tet*(X3) identified in *A. baumannii* 34AB (MK134375) and *A. indicus* AI2 (GCA012366935) are as the reference for the comparison of *tet*(X3); the **g**enomic context of *tet*(X6) identified in *P.* genomospecies 6 T60 (CP043925) and *A. indicus* CMG3-2 (CP044446) are as the reference for the comparison of *tet*(X6).

The genetic environment of *tet*(X6) was much more diverse than that of *tet*(X3) detected in our collection (Figure 2B). A 7,270-bp composite structure [ΔIS*CR2*-IS*30*-*tet*(X6)-*abh*-*guaA*-IS*CR2*] was detected in pAT232, which is similar with the prototype [ΔIS*CR2*-*tet*(X6)-*abh*-*guaA*-IS*CR2*] identified in pAT205 and a *Proteus* genomospecies 6 strain (26, 27), except for the insertion of an IS*30* (Figure 2B). The *tet*(X6) located within a 6,885-bp region [IS*CR2*-*fabF*-*tet*(X6)-*abh*-*glmM*-*sul2*] in pAT235 (Figure 2B), which shares 100% coverage and 99.58% identity with that detected on the chromosome of an *A. indicus* strain Q186-3_T and 100% coverage and 98.70% identity with pABF9692 carried by an *A. baumannii* strain (CP048828). In strain AT185, the genetic context of one copy of *tet*(X6) was identical to that detected in pAT235, and a truncated structure was found for the other copy (Figure 2B). The IS*CR2*-*fabF*-*tet*(X6)-*abh* fragment was also found on the chromosome of *A. indicus* strain LYS68A (CP070997) and *A. baumannii* strain 31FS3-2 (CP0445177), indicating that this structure might mediate the mobilization of *tet*(X6) on the plasmid and chromosome of *Acinetobacter* spp..

### A tet(X3)-carrying plasmid is self-transmissible from A. towneri to A. baumannii and increased the resistance to tetracyclines and growth rate

Conjugation assay was performed to test the transferability of *tet*(X)-encoding plasmids. We only obtained tigecycline-resistant transconjugants of *A. towneri* strain AT181 with frequencies at 1.85×10^-6^ per recipient cell. Multiple attempts of plasmid transfers failed when *E. coli* strain EC600 was used as a recipient. Compared with that of the recipient strain ATCC17978, the MIC value of tigecycline and the other tetracyclines against the transconjugant ATCC17978-pAT181 increased by 128-fold and 64∼512-fold, respectively (Table S2). To understand the transmission pattern of *tet*(X3) (i.e., by plasmid or by a circular form), WGS were performed for ATCC17978-pAT181 and ATCC17978 to detect the transferrable structure of *tet*(X3). A unique plasmid pAT181 was detected in the transconjugant ATCC17978-pAT181, demonstrating that the transmission of tigecycline resistance was mediated by pAT181 (Figure S2). This is different from another self-transmissible *tet*(X3)-harboring plasmid p10FS3-1-3 that the transfer of p10FS3-1-3 into *A. baylyi* ADP1 did not bring significant additive effect on the resistance to tetracyclines (20). To our best knowledge, this is the first report showing that the horizontal transfer of *tet*(X3)-carrying plasmid conferring tetracyclines resistance to the recipient.

*tet*(X3) was stable in the recipient strain ATCC17978 without antibiotic stress after 10-day passage, with 100% retention rate, indicating that pAT181 is able to be stably maintained in ATCC17978. The growth rate of the transconjugant ATCC17978-pAT181 increased compared with that of ATCC17978, and the doubling time shortened from 4.59 h to 2.91 h (Figure 3). These results suggest that pAT181 could facilitate the dissemination of *tet*(X3) among *Acinetobacter* spp..

**Figure 3.**
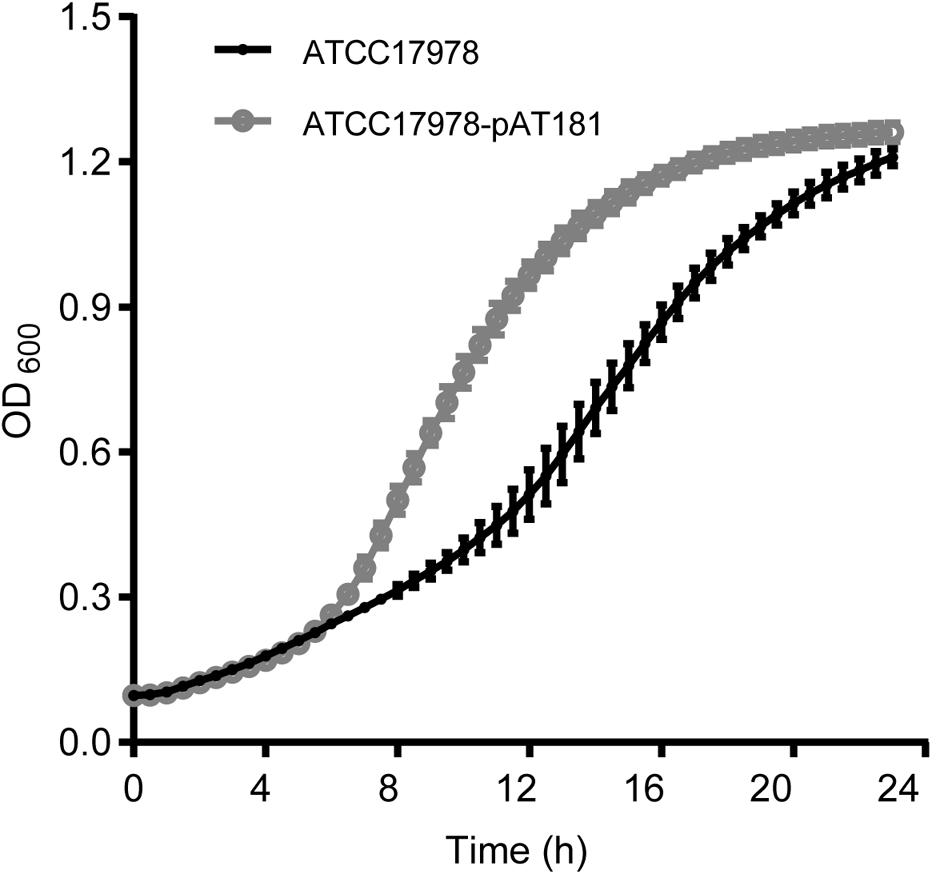
The growth curve of the recipient strain *A. baumannii* ATCC17978 and the transcongjutant ATCC17978-pAT181. The optical density at 600 nm was record every 30 min at 37°C. The assay was in triplicate.

### *tet*(X3) and *tet*(X6) are the prevalent alleles of *tet*(X) family and mainly sporadically disseminate in four species of *Acinetobacter* spp

As shown in this and other studies (7, 16, 17, 20, 23), *Acinetobacter* spp. is the major host of *tet*(X)s. To fully understand the distribution of *tet*(X)s among *Acinetobacter* spp., the nucleotide-acid sequences of 15 known *tet*(X) alleles and their variants were blasted against 10,680 *Acinetobacter* genomes retrieved from GenBank. *tet*(X3) was found in 249 strains; *tet*(X4) in 9 strains; *tet*(X5), *tet*(X5.2) and *tet*(X5.3) in 61 strains; *tet*(X6) in 53 strains; *tet*(X13), an one-residue variant of *tet*(X6), was found in 4 strains. These data reveal that *tet*(X3), *tet*(X5.2) and *tet*(X6) are the prevalent *tet*(X) genes among *Acinetobacter* spp..

Species identification showed three predominant *Acinetobacter* species carrying *tet*(X3), i.e. *A. indicus* (27.71%; 69/249), *Acinetobacter* sp002018365 (26.51%; 66/249) (an unnamed species with *Acinetobacter* sp. ANC 4845 as the reference) and *A. towneri* (13.65%; 34/249). Except for *A. variabilis* (11.32%; 6/53), *A. indicus* (22.64%; 12/53), *Acinetobacter* sp002018365 (20.75%; 11/53) and *A. towneri* (11.32%; 6/53) are also the predominant species carrying *tet*(X6). The species distribution of *tet*(X5.2) was similar with *tet*(X6), and the major species include *A. indicus* (22.64%; 12/53), *Acinetobacter* sp002018365 (20.75%; 11/53), and *A. towneri* (11.32%; 6/53), *A. variabilis* (11.32%; 6/53) and *A. lwoffii* (11.32%; 6/53). These results indicate that *A. indicus* and *Acinetobacter* sp002018365 are the most prevalent species carrying *tet*(X) genes.

To further evaluate the dissemination pattern of *tet*(X3) and *tet*(X6) among *Acinetobacter* population, we performed phylo-genomic analysis for *tet*(X3)/*tet*(X6)-positive strains of four major hosts as representatives, i.e. *A. indicus*, *Acinetobacter* sp002018365, *A. towneri* and *A. variabilis* (Figure 4; Figure S3). Most strains of each species shared a distant relationship, and no epidemic clones were detected. Two inter-regional transmission events were detected for 4 (no SNPs) and 5 (0-1 SNP) strains of *A. indicus,* and one cross-host event (pig and environment) was detected for 4 (1-44 SNPs) strains of *Acinetobacter* sp002018365 (Figure 4). The data suggest that *tet*(X3) and *tet*(X6) mainly sporadically disseminate among *Acinetobacter* population.

**Figure 4.**
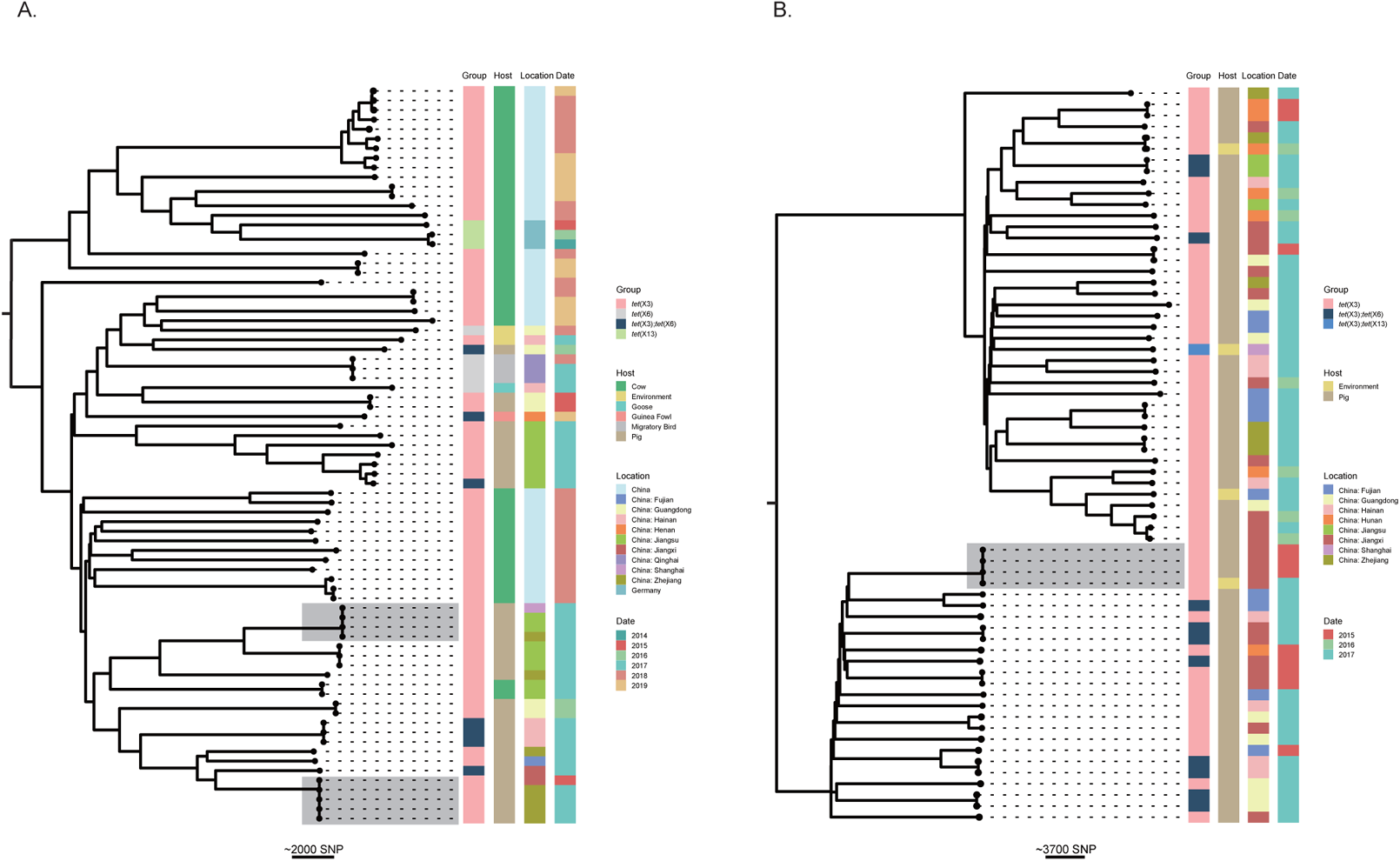
Phylogenetic analysis of *tet*(X3)/*tet*(X6)-encoding *A. indicus* (A) and *Acinetobacter* sp002018365 (B) genomes retrieved from GenBank. The core-genome SNPs of *tet*(X)-encoding strains were used to generate the phylogenetic tree. The tree is mid-point rooted. The *tet*(X) genes (group), isolate source (host), sampling location (location) and years (date) of strains are shown at the right side of the tree in different colors. Two inter-regional transmission events for 4 and 5 strains of *A. indicus*, and one cross-host event for 4 strains of *Acinetobacter* sp002018365 are highlighted by shading. The scale bar represents the number of SNPs.

### The structures of *tet*(X3)/*tet*(X6) plasmidome are highly diverse and no epidemic plasmids have emerged among *Acinetobacter* population yet

To explore the role of plasmids in the disseminations of *tet*(X3) and *tet*(X6) in *Acinetobacter* spp., we here intended to dissect the genetic relatedness of *tet*(X3) and *tet*(X6)-harboring plasmids. Four circularized *tet*(X3)/*tet*(X6)-harboring plasmids obtained in this study and all finished *tet*(X3)/*tet*(X6)-harboring plasmids deposited in GenBank [n = 30; 18 for *tet*(X3), 6 for *tet*(X6), and 6 for *tet*(X3) and *tet*(X6)] were analyzed at first. All but two of these publicly available plasmids were collected between 2009 and 2020 in China, and 25 were identified in *Acinitobacter* spp. (Figure 5). Pairwise comparisons using nucleotide-acid sequences revealed that most of the 26 *tet*(X3)-harboring plasmids (including the 6 *tet*(X3)-*tet*(X6)-harboring plasmids) share a coverage lower than 65%, indicating a highly diverse structure for the plasmidome of *tet*(X3) (Figure 5A). Four of the 6 *tet*(X3)-*tet*(X6)-positive plasmids share a high similarity (>89.83% coverage; >85% identity), suggesting that they were derived from an ancestor. The 4 plasmids were hosted in *A. schindleri* and *A. indicus* isolated from goose and soil collected in different provinces of China (Figure 5A), indicating that cross-species, cross-sector (poultry and environment) and/or cross-region transmission has occurred for these plasmids. A similar transmission event was observed for another three *tet*(X3)-encoding plasmids (pAT181, pAT184 and p10FS3-1-3) carried by *A. towneri* and a novel species of *Acinetobacter* as aforementioned (Figure 5A).

**Figure 5.**
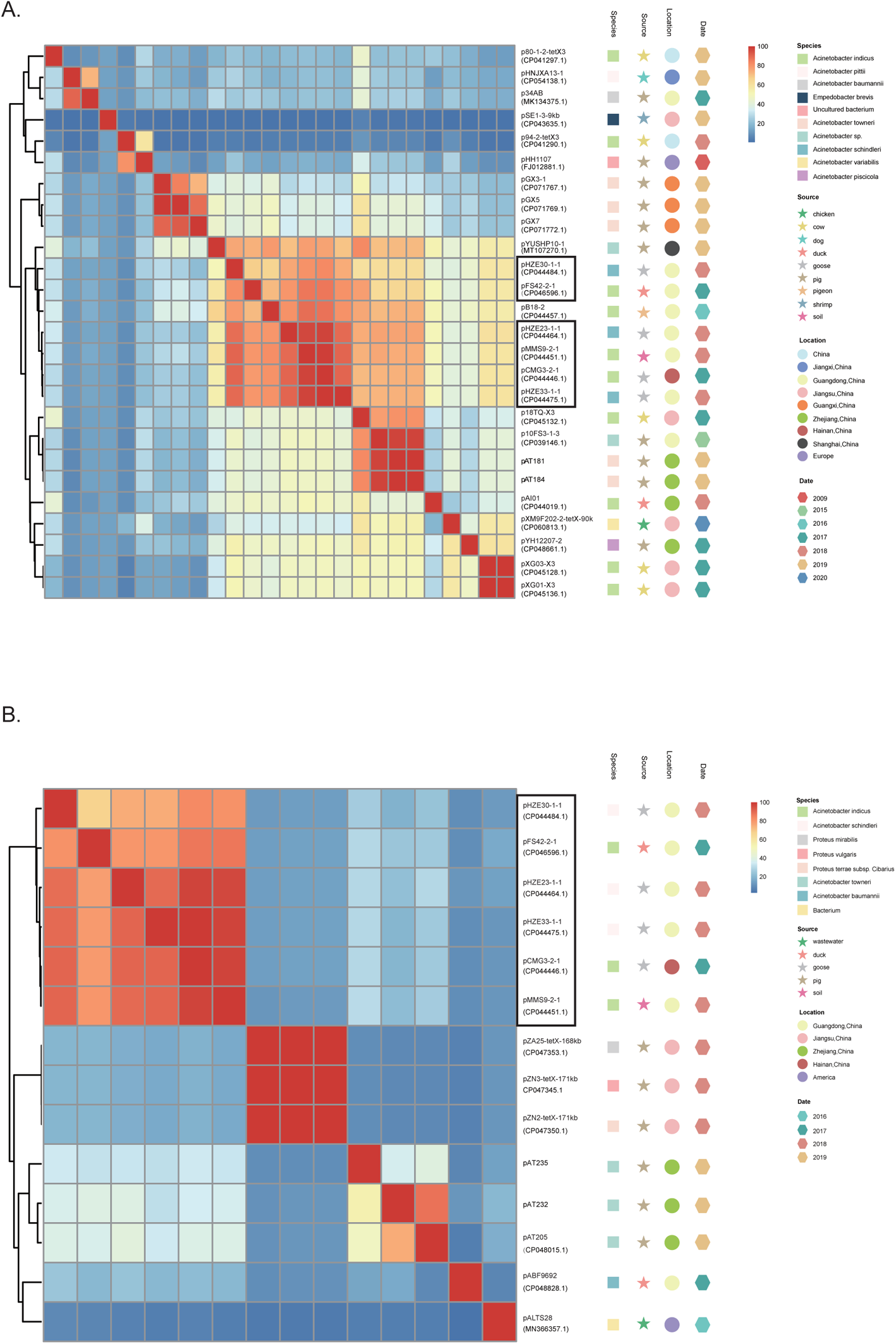
Pairwise sequence comparisons between circularized *tet*(X3)/*tet*(X6)-carrying plasmids. The heat map shows the percentage of aligned bases between pairs of *tet*(X3)-carrying plasmids **(A)** and *tet*(X6)-carrying plasmids **(B)**. The row and column orders are the same. The information of host species, sampling source, sampling location and isolation years are indicated by colored graphic subsequent the phylogenetic tree. The 6 plasmids co-harbored *tet*(X3) and *tet*(X6) genes are boxed.

The pairwise sequence comparison of 14 *tet*(X6)-harboring plasmids (including the 6 *tet*(X3)-*tet*(X6)-harboring plasmids) showed that, the 5 *tet*(X6)-harboring plasmids carried by *Acinetobacter* and an unknown species share a low similarity, except for pAT232 and pAT205 as aforementioned (Figure 5B). They are different from the 3 *tet*(X6)-harboring plasmids (pAZ25, pZN3 and pZN2) carried by *Proteus* species, and the 6 *tet*(X3)-*tet*(X6)-harboring plasmids (Figure 5B). This suggests that the *tet*(X3)-*tet*(X6)-harboring plasmids might be resulted from the capture of *tet*(X6) by *tet*(X3)-harboring plasmids.

To further understand the distribution of *tet*(X3)-harboring plasmids among *Acinetobacter* spp., we selected 17 plasmids out of 26 *tet*(X3)-harboring plasmids as reference according to their similarities (< 80% coverage and identity). The 17 plasmids were blasted against the 243 *tet*(X3)-positive genomes (6 genomes with chromosome-encoding *tet*(X3) were excluded), and no epidemic plasmids were found (Figure 6A). To evaluate structural conservation of plasmids amongst *tet*(X3)-positive isolates, we mapped the 243 genomic sequences against the 17 representative plasmid sequences (Fig. 6B). This revealed that plasmid structures were highly diverse amongst isolates (mean plasmid coverage range 12.09-55.05%). Using a cutoff range of >80% coverage and >90% identity, we found that a pGX5-like plasmid was hosted in 36 strains belonging to different species (20 *A. towneri* strains, 10 *A. variabilis* strains, 4 *Acinetobacter* sp002018365 strains and 2 *A. indicus* strains), and a p34AB-like, a p94-2-*tet*X3-like, a pXM9F202-2-*tet*X-90k-like and a p10FS3-1-3-like plasmid were found in 17, 9, 8, and 7 strains belonging to different species, respectively (Figure 6A). These data suggest that the current dissemination of *tet*(X3) in *Acinetobacter* is mainly mediated by various plasmids, and cross-species transmissions mediated by few of them might have occurred in a small proportion of strains.

**Figure 6.**
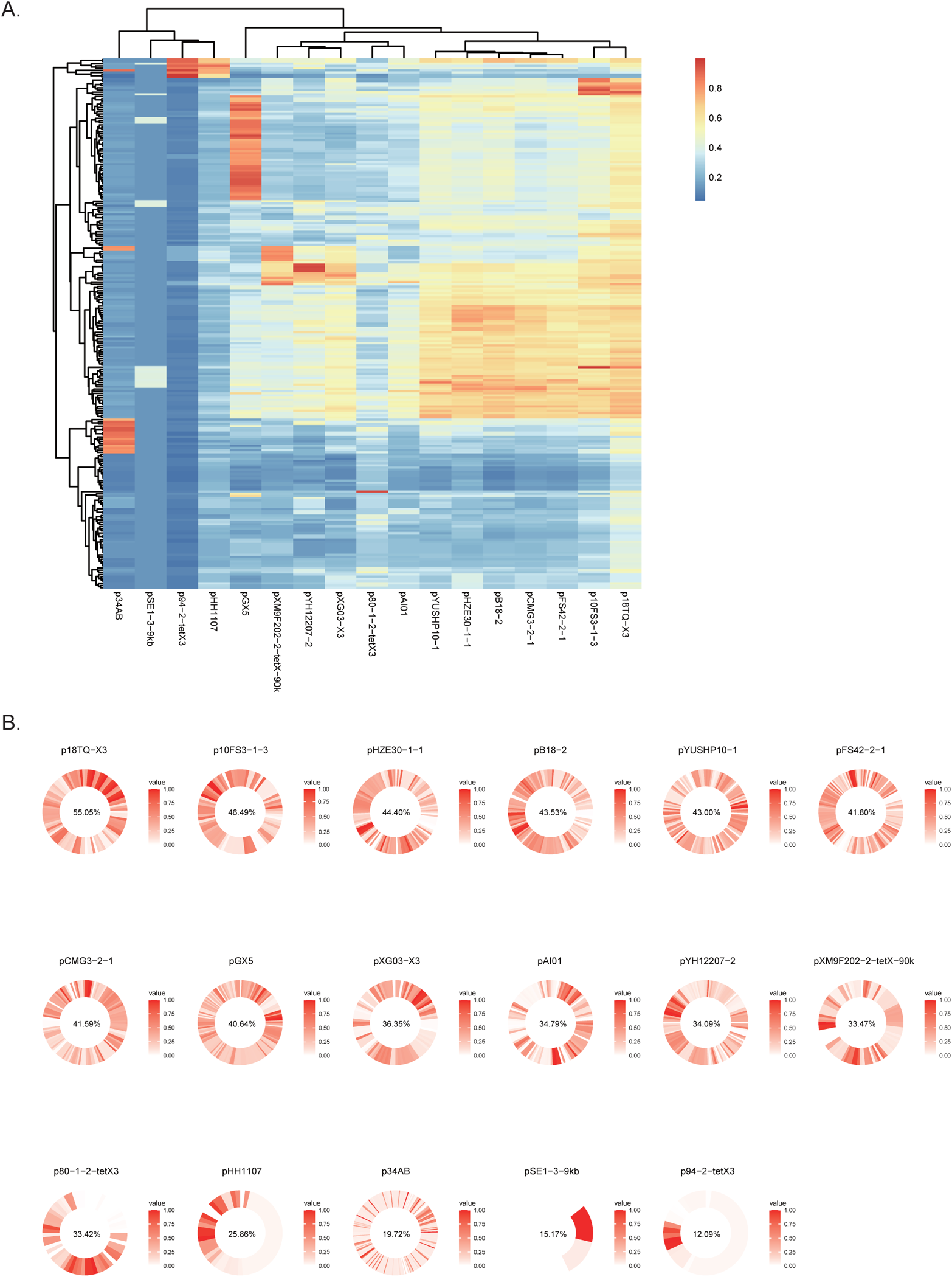
Analysis of *tet*(X3) Plasmidome. (A) Blasting results of the 17 representative *tet*(X3)-carrying plasmids against 243 *tet*(X3)-positive genomes. The heat map shows the percentage of aligned bases between pairs of *tet*(X3)-positive plasmids and genomes. (B) Conservation of reference plasmid genes amongst 243 genome sequences of *tet*(X3)-carrying *Acinetobacter* spp.. The frequency of each gene in the reference plasmid is shown in circularized heatmaps. Genes order in the corresponding reference plasmid are around the cell. The mean coverage of the reference plasmids sequence is indicated in percentages after the plasmid name.

### The promoter of *tet*(X3) and *tet*(X6) is interchangeable

Our previous study showed that *tet*(X3) confers higher tigecycline resistance than *tet*(X6) (26). In order to understand whether this difference is resulted from the different promoters of the two genes, constructions of promoter-exchanged overexpression plasmids were performed. Transformants carrying the expression cassette promoter*_tet_*_(X6)_-ORF*_tet_*_(X3)_ [*tet*(X3) ORF followed *tet*(X6) promoter] exhibited the same level of resistance with that of the original cassette promoter*_tet_*_(X3)_-ORF*_tet_*_(X3)_ (Table 3). Likewise, the reconstruction of expression cassette promoter*_tet_*_(X3)_-ORF*_tet_*_(X6)_ did not alter the activity of *tet*(X6) either (Table 3). These results suggest that the tigecycline resistance activity of *tet*(X3) and *tet*(X6) could be determined by the sequence of ORFs and the promoter of *tet*(X3) and *tet*(X6) is interchangeable.

**Table 3.** MIC values of tetracyclines tested in this study

| Strains | MIC (mg/L) |  |  |  |  |  |  |  |
| --- | --- | --- | --- | --- | --- | --- | --- | --- |
|  | TGC | TET | OTC | CTC | DMC | DOX | MIN | ERV |
| **DH5α-*tet*(X3)promoter-*tet*(X6)ORF** | 8 | >128 | >128 | >128 | >128 | >128 | 8 | 8 |
| **DH5α-*tet*(X6)promoter-*tet*(X6)ORF** | 8 | >128 | >128 | >128 | >128 | >128 | 16 | 8 |
| **DH5α-*tet*(X6)promoter-*tet*(X3)ORF** | 32 | >128 | >128 | >128 | >128 | >128 | 64 | 32 |
| **DH5α-*tet*(X3)promoter-*tet*(X3)ORF** | 16 | >128 | >128 | >128 | >128 | >128 | 32 | 32 |

Abbreviation: TGC, Tigecycline; TET, Tetracycline; OTC, Oxytetracycline; CTC, Chlortetracycline; DMC, Demeclocycline; DOX, Doxycycline; MIN, Minocycline; ERV, Eravacycline.

## Discussion

A total of 15 *tet*(X) alleles have been reported since 1990 (9), and they have spread to cover 5 of the 7 continents (8). Recent surveillance for *tet*(X)s reveals the wide range of ecosystems, including soil, sewage, animals, hospitals, livestock farms and human gut (14–16, 28). The *tet*(X)-positive isolates are especially prevalent in livestock and poultry, like pigs, cows, chicken, and less in shrimp, migratory birds and waterfowls (7, 16, 18, 23, 28–31). In this study, we comprehensively characterized *tet*(X)-positive strains collected from different livestock farms (swine farms, dairy farms and sheep farms), and we found that *tet*(X)-positive strains were exclusively isolated from swine farms (Table 1). A similar finding has been reported recently that *tet*(X3)-positive *Acinetobacter* spp. isolates were exclusively detected in the intensive pig farms in China (20). These results suggest that the dissemination risk of *tet*(X) genes to human from pigs could be much higher than from other livestock.

Current surveillances show that *Acinetobacter* spp. is the major reservoir of *tet*(X) genes (17, 20, 23). In this study, *A. towneri* was found to be the major host of *tet*(X3) and *tet*(X6). A recent surveillance of *tet*(X)-positive *Acinetobacter* isolates from human, animal, and their surrounding environments conducted between 2015 and 2018 shows that *A. towneri* and *A. indicus* following a novel species of *Acinetobacter* were the major hosts of *tet*(X3), *tet*(X4) and *tet*(X5) (20). This indicates that the diversity of *tet*(X) hosts may be source- and/or geographic-dependent. To fully understand the distribution of *tet*(X)s in *Acinetobacter* population, we searched 15 *tet*(X) alleles and their variants in all *Acinetobacter* genomes available in GenBank, and results revealed that *tet*(X3) and *tet*(X6) are the predominant alleles mostly associated with livestock, and *A. towneri* is the third prevalent species carrying *tet*(X3) and *tet*(X6) following *A. indicus* and *Acinetobacter* sp002018365. Further analysis showed that the population structure of the four major species is highly diverse (Figure 4 and S3), suggesting that *tet*(X3) and *tet*(X6) are mainly sporadic dissemination. However, few inter-regional transmission events were detected here, highlighting the needs for controlling the dissemination of *tet*(X3) and *tet*(X6) positive *Acinetobacter* spp., especially from livestock to humans.

First identification of plasmid-borne *tet*(X3) and *tet*(X4) causing the horizontal transfer of tigecycline resistance has highly aroused the public attention. Since then, numerous *tet*(X) alleles have been continuously identified either on chromosomes or on plasmids in various bacterial species. However, whether plasmids are the major vectors of plasmid-borne *tet*(X)s remains unclear. Pioneer studies have shown the importance of IS*CR2*-mediated *tet*(X) transposition structure (7, 17). The rolling-circle transposition has been experimentally confirmed by using the cassette “Δ*tpnF*-*tet*(X3)-hp-hp-IS*CR2*” clone, and inverse PCR assays identified “IS*CR2*-*xerD*-*tet*(X3)-*res*-ORF1” and “IS*CR2*-ORF2-*abh*-*tet*(X4)” minicircles in different studies (7, 20). In our study, IS*CR2* was found upstream or downstream of *tet*(X3) and *tet*(X6) genes. Albeit we did not test the transferability of the IS*CR2*-mediated *tet*(X) transposition structure, the genetic context of *tet*(X3) carried by 249 genomes of *Acinetobacter* species were comprehensively compared. The proportion of the structure IS*CR2*-*xerD*-*tet*(X3)-*res*-IS*CR2* might be up to 86.35% (215/249), implying the critical role of IS*CR2* in the dissemination of *tet*(X3).

Of note, we found that a *tet*(X3)-encoding plasmid pAT181 was self-transmissible from *A. towneri* to *A. baumannii*, and conferred tetracyclines resistance to the recipient. Currently, very few studies have identified self-transmissible plasmids carrying *tet*(X)s. Chen *et al*. reported the conjugability of a *tet*(X3)- and *tet*(X5.3)-harboring plasmid pYH12207-2 from *A. piscicola* to *A. baylyi* ADP1, and the conjugability of a *tet*(X3)-harboring plasmid p10FS3-1-3 from an *Acinetobacter* novel species to *A. baylyi* ADP1. However, these two plasmids did not enhance the resistance to tetracyclines in the recipient strain (20). This is different from our findings that the transfer of pAT181 to the recipient resulted in a 64-512-fold increase of tetracyclines resistance (Table S2). Remarkably, the donor strain of pAT181 is a tigecycline-susceptible *A. towneri* strain, and the recipient strain is *A. baumannii*, suggesting that the expression of *tet*(X3) could be species-dependent. More than half of *tet*(X3)/*tet*(X6)-positive *A. towneri* strains were tigecycline-susceptible in this study, indicating the silent transmission of *tet*(X3)/*tet*(X6) in *A. towneri.* Concerningly, pAT181 with a relatively high transfer frequency (10^−6^) did not impose fitness cost but increased the growth rate of the recipient. It is suggested that successful disseminations of resistance plasmids largely depends on the fitness cost imposed on hosts (32). No fitness cost imposed on hosts by obtaining pAT181-like plasmids would greatly facilitate their spread, thus may contribute to the propagation of *tet*(X3) gene in the future. Additionally, although no epidemic plasmids of *tet*(X3) have been detected currently, several plasmids were found circulating in a small proportion of strains. It is possible that these plasmids could become epidemic after transmitting to other hosts in the future.

## Conclusions

Our study evidence that the predominate *tet*(X) alleles, *tet*(X3) and *tet*(X6), disseminate sporadically in *Acinetobacter* population. Currently, the dissemination of *tet*(X3) and *tet*(X6) is mainly limited among livestock-associated sites. Continuous surveillance for *tet*(X)s in the context of One Health is necessary to prevent them from transmitting to humans.

## Materials and Methods

### Screenings of *tet*(X)-positive *Acinetobacter* spp. strains

Five hundred and thirty-four non-repetitive fecal samples were collected from 6 livestock farms locating in Zhejiang province in 2019, including 2 swine farms, 2 dairy farms and 2 sheep farms. Environmental samples were collected from soil (n=72) and water (n=78) surrounding the farms in parallel. These samples were initially enriched in LB medium (5 g/L yeast extract, 10 g/L tryptone, 10 g/L NaCl) for 6 hours and spread on CHROMagar^TM^ *Acinetobacter* medium plates (CHROMagar, Paris, France) to recover *Acinetobacter* spp. strains. PCR screens of *tet*(X) alleles were performed as previously described (26).

### Antimicrobial susceptibility testing (AST)

The minimum inhibitory concentration (MICs) for all *tet*(X) positive strains were determined using microbroth dilution method according to the guideline of Clinical and Laboratory Standards Institute (CLSI) (29th edition) (33). The tested drugs included tigecycline, tetracycline, eravacycline, minocycline, doxycycline, demeclocycline, chlortetracycline, oxytetracycline, colistin, cefoperazone-sulbactam, trimethoprim-sulfamethoxazole, gentamicin, amikacin, levofloxacin, ciprofloxacin, meropenem, cefepime, ceftriaxone, and ceftazidime. The breakpoint for tetracycline was interpreted as ≥ 16 mg/L for *Acinetobacter* spp., *Enterobacteriacea* and non-*Enterobacteriaceae* according to CLSI (33). The breakpoint for tigecycline and eravacycline was interpreted as > 0.5 mg/L for *Enterobacteriaceae* according to EUCAST V10 (34). *E. coli* ATCC25922 was used as the quality control strain.

### Whole genome sequencing (WGS) and bioinformatic analysis

Genomic DNAs of *tet*(X)-positive isolates were extracted by using Puregene Yeast/Bact Kit B (Qiagen, Gaithersburg, MD, Germany) according to the instruction of the manufacture, and were sequenced by using Hiseq 4000 system (Illumina, San Diego, United States). The average nucleotide identity (ANI) was calculated by using FastANI (35). Sequence similarity of *tet*(X)-harboring plasmids was analyzed by using BRIG v0.95 (36). Representative strains with various genetic context of *tet*(X) genes were selected out to be further sequenced by using PromethION platform (Nanopore, Oxford, UK). Hybrid assembly of short reads and long reads sequencing data was performed by using Unicycler version 0.4.8 (37).

Phylogenetic analysis was performed by using Parsnp v1.2 (38), and the number of SNPs (single nucleotide polymorphisms) among the core genomes were determined by MEGA X (39). Functional annotation was performed using RAST server (40). Antibiotic resistance genes (ARGs) were identified by using ResFinder 4.0 (41) and CARD (https://card.mcmaster.ca/) with the threshold of nucleotide-acid identity >90% and coverage >90%. Synteny analysis was performed by using Easyfig (42).

### Compilation of genomic data set and plasmidome analysis

All assembled genomes of *Acinetobacter* spp. (n= 10,680) deposited in GenBank (as of 31th May 2021) were downloaded to search *tet*(X) alleles. The fifteen *tet*(X) alleles were queried in these genomes by blasting against their nucleotide-acid sequences using a cutoff as 99% identity and 100% coverage.

Conservation of reference plasmid genes was calculated as previously described (43). Briefly, RedDog pipeline (https://github.com/katholt/RedDog) was used to simulate 100-bp reads from *tet*(X3)-carrying genomes. To calculate the coverage of each representative plasmid in each genome, those 100-bp reads were mapped against representative *tet*(X3)-harboring plasmids by using Bowtie2 v2.2.9 (44). The proportion of *tet*(X3)-carrying genomes containing annotated genes of each reference plasmid was calculated according to the gene presence/absence table reported by Red-Dog (at least five reads covering ≥ 95% of the length of the gene was defined as presence) and plotted as circular heatmaps using ggplot2 in R (geom_tile for heatmap grid and coord_polar for circularise).

Pairwise sequence comparison of circularized plasmids was performed as previously described (45). Briefly, the length of nucleotide-acid sequence that could be aligned between pairs of plasmids and the number of SNPs among the aligned regions were determined by NUCmer v3.1 (46) from the MUMmer package. The percentage of aligned bases between pairs of complete plasmids was showed in heatmap generated by the “gplots” package (v3.1.1) in R v4.0.5 (https://www.r-project.org/).

### Conjugation assay

The transmissibility of *tet*(X3) and *tet*(X6) was evaluated by conjugation assay. Briefly, *tet*(X)-carrying *Acinetobacter* strain as a donor strain was mixed with rifampicin-resistant *A. baumannii* ATCC17978 or rifampicin-resistant *E. coli* EC600 as a recipient strain at the ratio of 1:1 by conjugational mating at 37°C without shaking for overnight. The transconjugants were selected on LB agar plates containing rifampicin (600 mg/L) and tigecycline (2 mg/L). The species of all putative transconjugants were verified by using MALDI-TOF mass spectrometry (Hexin, Guangzhou, China). PCR verifications of *tet*(X) genes were performed for the putative transconjugants of which the species was confirmed as *A. baumannii* or *E. coli*. Transfer frequency was calculated as the number of transconjugants obtained per donor. Growth of donor strain and transconjugants were measured by determining the optical density at 600 nm (OD_600_) every 30 min.

### Plasmid stability testing

Plasmid stability was estimated according to a previous study with minor modifications (47). Transconjugants were cultured in antibiotic-free LB broth at 37°C for 24 h. The 24h-growth cultures were diluted with the ratio of 1:100 in fresh LB medium. These freshly inoculated cultures constituted time point zero, and cultures were grew at 37°C in a shaking bath (200 rpm) and went on serial passages for 10 days (approximately 200 generations). Cultures were diluted and plated onto antibiotic-free LB plates every 24 h. The colonies growing on antibiotic-free LB agar plates were randomly selected (∼50 colony per day) for *tet*(X)-specific PCRs to determine the proportion of *tet*(X)-positive bacteria in each population. Plasmids were considered stable when the retention rates were still over 80% at the end of the experiment. The plasmid stability was evaluated in triplicate.

### Functional cloning

Predicted promoters of *tet*(X3) and *tet*(X6) according to softberry (http://www.softberry.com/berry.phtml?topic=bprom&group=programs&subgroup=gfindb) were fused with open reading frames (ORFs) of *tet*(X3) and *tet*(X6) to construct promoter exchanged clones, respectively. Briefly, the promoter region of *tet*(X3) was amplified using primers PstI-*tet*(X3)-F-P (5’-cgctgcagTACCACCAAGGGAATGGAAC-3’) and X3P+X6O-R (5’-GTTCGCTGGTTTTAATGTCAATCAAAAATGGCACATAACAAG-3’), and the ORF of *tet*(X6) was amplified using primers X3P+X6O-F (5’-CTTGTTATGTGCCATTTTTGATTGACATTAAAACCAGCGAAC-3’) and XbaI-*tet*(X6)-R (5’-cgtctagaTTTCTCTTTCATTTCCTCGCC-3’). The derived amplicons were fused by the fusion PCR using primers PstI-*tet*(X3)-F-P and XbaI-*tet*(X6)-R, resulting in an X3P-X6O fragment. The digested X3P-X6O fragment was cloned into pUC19 to construct pUC19-X3P-X6O. Likewise, primers PstI-*tet*(X6)-F-P (5’-cgctgcagATGGTTGCAGACCTTGACGA-3’) and X6P+X3O-R (5’-CGTATCTATTCGCATTGTCATCTAATGTCTGTCAATTTAATC-3’) were used to amplify the promoter region of *tet*(X6). Primers X6P+X3O-F (5’-GATTAAATTGACAGACATTAGATGACAATGCGAATAGATACG-3’) and XbaI-*tet*(X3)-R (5’-cgtctagaGCAAAACTGCTTGTTAGTAGC-3’) were used to amplify the ORF of *tet*(X3).The fused amplicon X6P-X3O was ligated into pUC19 to construct pUC19-X6P-X3O. The recombinant plasmid was transformed into *E. coli* DH5*α* competent cells by heat shock. Transformants were selected on LB agar plates containing 100 mg/L ampicillin. *tet*(X3) and *tet*(X6) with parental promoters were individually cloned into pUC19 as positive controls.

## Statistical analysis

Statistical analysis was performed using unpaired *t*-test analysis, and statistical significance is taken as *p* < 0.05.

## Ethics approval and consent to participate

Not applicable.

## Consent for publication

Not applicable.

## Data availability

The genome sequences of *tet*(X) positive strains have been submitted to GenBank, and the accession numbers are listed in Table 1.

## Authors’ contributions

KZ, Y-YC, Y-HX, and R-CC designed the study. Y-YC, YC, and F-MH collected the data. Y-YC and YL analyzed and interpreted the data. Y-YC and KZ wrote and revised the manuscript. All authors reviewed, revised, and approved the final report

## Acknowledgements

This work was supported by the National Key Research and Development Program of China (grant number 2017YFC1200200); the National Natural Science Foundation of China (grant numbers 81902029); Shenzhen Basic Research Key projects (grant numbers JCYJ20200109144220704) and Shenzhen Basic Research projects (grant numbers JCYJ20190807144409307; JCYJ20190807150401657).

## Figure legends

**Supplementary figure 1. Comparative analysis of *tet*(X)-encoding plasmids in this study. (A)** The inner ring represents the circularized *tet*(X3)-encoding plasmid pAT181 as reference; **(B)** The inner ring represents the circularized *tet*(X6)-encoding plasmid pAT232 as reference; **(C)** The inner ring represents the circularized *tet*(X6)-encoding plasmid pAT235 as reference; **(D)** The inner ring represents the circularized *tet*(X6)-encoding plasmid pAT205 as reference.

**Supplementary figure 2. Verification of horizontal transfer of *tet*(X3)-encoding plasmid pAT181 in the transconjugant ATCC17978-pAT181.** The inner ring represents pAT181 as reference. The outer ring represents the mapping result of exogenous DNA in the tansconjugant AB181 compared with the recipient strain ATCC17978.

**Supplementary figure 3. Phylogenetic analysis of *tet*(X3)/*tet*(X6)-encoding *A. towneri* (A) and *A. variabilis* (B) genomes retrieved from GenBank.** The core-genome SNPs of *tet*(X)-encoding strains were used to generate the phylogenetic tree. The tree is mid-point rooted. The *tet*(X) genes (group), isolate source (host), sampling location (location) and years (date) of strains are exhibited at the right side of phylogenetic tree in different colors.

